# Sex-specific single cell-level transcriptomic signatures of Rett syndrome disease progression

**DOI:** 10.1101/2024.05.16.594595

**Authors:** Osman Sharifi, Viktoria Haghani, Kari E. Neier, Keith J. Fraga, Ian Korf, Sophia M. Hakam, Gerald Quon, Nelson Johansen, Dag H. Yasui, Janine M. LaSalle

## Abstract

Dominant X-linked diseases are uncommon due to female X chromosome inactivation (XCI). While random XCI usually protects females against X-linked mutations, Rett syndrome (RTT) is a female neurodevelopmental disorder caused by heterozygous *MECP2* mutation. After 6-18 months of typical neurodevelopment, RTT girls undergo poorly understood regression. We performed longitudinal snRNA-seq on cerebral cortex in a construct-relevant *Mecp2e1* mutant mouse model of RTT, revealing transcriptional effects of cell type, mosaicism, and sex on progressive disease phenotypes. Across cell types, we observed sex differences in the number of differentially expressed genes (DEGs) with 6x more DEGs in mutant females than males. Unlike males, female DEGs emerged prior to symptoms, were enriched for homeostatic gene pathways in distinct cell types over time, and correlated with disease phenotypes and human RTT cortical cell transcriptomes. Non-cell-autonomous effects were prominent and dynamic across disease progression of *Mecp2e1* mutant females, indicating wild-type-expressing cells normalizing transcriptional homeostasis. These results improve understanding of RTT progression and treatment.

## Introduction

Rett syndrome is a neurodevelopmental disorder primarily affecting females and is characterized by a range of symptoms such as loss of speech, motor abnormalities, and developmental regression at about 6-18 months of age^1^. Rett syndrome most often occurs through spontaneous germline mutations in the X linked gene *MECP2* that are mostly missense or truncation mutations^2^. *MECP2* encodes the DNA binding protein, methyl CpG binding protein 2 (MeCP2), which is a critical regulator of neuronal gene expression in the brain^3^. Among the two alternatively spliced *MECP2* transcripts, only the MeCP2e1 isoform contributes to RTT disease phenotypes^4^. However, most mouse studies of RTT utilize the exon 3-4 knockout model in *Mecp2*^*-/*y^ males^5^, which is a null model effective for studying MeCP2 function, but not a construct- or sex-relevant model for human RTT. RTT females are heterozygous for *MECP2* (*MECP2*^*-/+*^) mutations and are therefore mosaic for both *MECP2* wild-type and mutant cells in brain. Prior studies in Rett syndrome have suggested potential non-cell-autonomous effects of MeCP2 deficiency on wild-type expressing cells in brain, but these effects have been poorly characterized at a cellular and molecular level^6–8^. RTT is characterized by a seemingly typical development in infancy, followed by progressive stages of regression in developmental milestones beginning around 6-18 months of age and lasting through early adulthood^8^. We have previously demonstrated that the *Mecp2e1* deficient mouse model of RTT, modeled after a human mutation, recapitulates the RTT-relevant extended period of disease symptom progression^4,9^. However, it is not known when and in which cell types the molecular changes responsible for disease progression occur in *MECP2* mutant females versus males.

To explore the effects of cellular mosaicism, sex, and cell type on the progression of disease in Rett syndrome, we employed single nuclei RNA-seq (sn-RNA-seq 5’) analysis in the cerebral cortex of the *Mecp2e1* mutant mouse model. We examined the influence of sex, cell type, cellular mosaicism, and disease stage, correlated with progressive disease phenotypes using a systems-level perspective. These results demonstrate that MeCP2 deficiency in females results shows an inherently different disease progression at the cellular and molecular level compared to males, involving non-cell-autonomous transcriptional changes to homeostatic gene pathways that correlate with disease phenotypes and stages.

## Results

### Experimental design to test longitudinal, cellular, and sex-specific transcriptional dysregulation in a symptomatically progressive mouse model of Rett syndrome

To identify sex, cell type, and disease stage specific transcriptional differences in *Mecp2e1* deficient mouse cortex, single nuclei RNA sequencing (sn-RNA seq 5’) analysis was performed to include the engineered mutation at the 5’ translational start site of the *Mecp2e1* isoform^4^. Three longitudinal post-natal time points were chosen to correspond to pre-symptomatic (PND 30), disease onset (PND 60) and late disease stages (PND 120 for *Mecp2e1*^*-/y*^ males, PND 150 for *Mecp2e1*^*-/+*^ females) compared to sex-matched wild-type (WT) littermates^9–11^ **(Figure 1a**). Cortical nuclei were assigned to 14 different cell types based on 3,000 cell marker genes from the Allen brain atlas cortex transcriptomics data^12^. 93,798 cells from both sexes, four genotypes and three time points were all clustered unsupervised (**Figure 1b**). Four excitatory neuron cell types were identified, corresponding with cortical layers 2 to 6 (L2-6), as well as six inhibitory cell types (Pvalb, Vip, Sst, Sncg, Lamp5) and four non-neuronal cell types (pericytes, endothelial, oligodendrocytes, astrocytes, non-neuronal including microglia). Unbiased marker genes for all 14 cell types were identified, supporting the distinction of our candidate cell types (**Figure 1c**).

**Figure 1.**
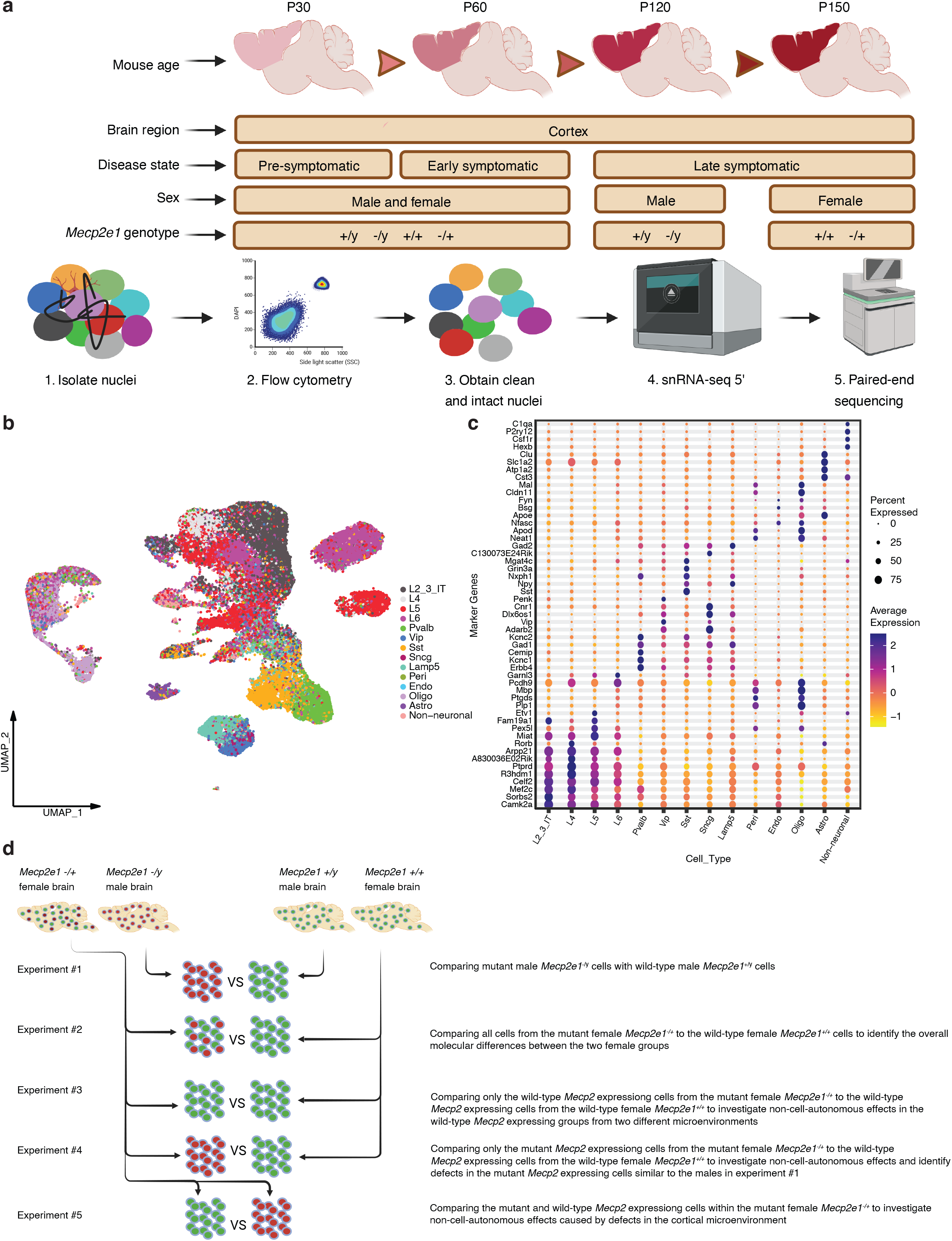
A scheme showing the overall mouse study design. **a**. Cortical samples were collected from postnatal mice at four different timepoints corresponding to three different disease stages (n = 28). Four different Mecp2e1 genotypes were considered that include both sexes. **B**. UMAP of the unsupervised clustering of cell types (n = 93,798 cells post QC) identified. Cell type labels were transferred from^38^ *Van, Yao et al. 2021*. **c**. Top gene markers for each cell type are shown on y-axis. The color refers to the average expression of genes in a cell type and the percent expressed describes the percentage of cells within a cell type that express each gene marker. **d**. Design of computational experiments comparing mutant to WT cells from mice of both sexes. Experiments 3 to 5 are comparing subtypes of cells in females due to X chromosome inactivation to examine potential non-cell-autonomous effects of *Mecp2e1* mutation.

Five separate hypotheses were tested, comparing cells across different genotypes and expression phenotypes (mutant vs wild-type-expressing cells within females). In addition to comparing cells from *Mecp2e1*^*-/y*^ to *Mecp2e1*^*+/y*^ (experiment 1) and *Mecp2e1*^*-/+*^ to *Mecp2e1*^*+/+*^ (experiment 2), wild-type *Mecp2e1* expressing cells from the *Mecp2e1*^*-/+*^ females were compared to the wild-type expressing cells from the *Mecp2e1*^*+/+*^ (experiment 3) and mutant *Mecp2e1* expressing cells from the *Mecp2e1*^*-/+*^ females were compared to either wild-type expressing cells from the *Mecp2e1*^*+/+*^ (experiment 4) or wild-type expressing cells within *Mecp2e1*^*-/+*^ females (experiment 5) to test for cell non-autonomous effects (**Figure 1d**).

## Sexually dimorphic trajectories of transcriptional dysregulation across cortical cell types

To accurately characterize alterations in gene transcript abundance, four computational methods for identifying differentially expressed genes (DEGs) from single nucleus RNA sequencing (snRNA-seq) data were evaluated with single cell data sets (Limma-VoomCC, Limma, EdgeR, and DESeq2) with partial overlap (**Supplemental Figure 1**). Ultimately, Limma-Voom Consensus Correlation (Limma-VoomCC) was selected for DEG analysis based on the ability to reveal high expressing DEGs amongst diverse gene transcripts expressed^13^. Further, Limma-VoomCC controlled for the inter-correlations of cells from the same animals^14,15^. Overall, Limma-VoomCC analyses of all cell types in experiments 1 and 2 revealed a total of 1436 significant DEGs after adjusting for false discovery (**Supplemental Table 1**). In males from experiment 1, 169 or 85% showed higher and 30 or 15% showed lower transcript levels in *Mecp2e1* mutant cortical cells compared to wild-type controls across the three time points, with fold changes ranging from a low of -1.99 for *Sst* to a high of +2.31 for *Cst3*. In females from experiment 2, 282 or 22% showed higher and 959 or 77% showed lower transcript levels in *Mecp2e1* mutant cortical cells compared to wild-type controls across the three time points, with fold changes ranging from a low of -2.69 for *Snhg11* to a high of +3.47 for *Ay036118* (**Supplemental Table 1**). DEsingle was also used as a complementary approach to identify lower confidence DEGs for transcripts expressed at low levels (**Supplemental Figure 2**). To ensure that DEGs detected were not due to changes in cell types, we examined cell proportions which did not show changes over time (**Supplemental Figure 3**). Cell clustering based on cell type, time point, sex and *Mecp2e1* genotype did not show evidence of batch effects (**Supplemental Figure 4a-d**). Futher, an analysis of the top high and low expressing genes showed that brain samples from replicate mice were consistent (**Supplemental Figure 5**).

Analysis of DEGs by both Limma-VoomCC and DEsingle revealed that cell type transcriptional changes associated with *Mecp2e1* deficiency were markedly different by sex and disease stage in multiple cortical cell types (**Figure 2, Supplemental Figure 2**). At the pre-symptomatic stage, *Mecp2e1*^*-/y*^ male P30 from experiment 1 cortical cells had only 9 DEGs compared with wild-type, including 3 DEGs in L2/3 neurons, 4 DEGs in L4 neurons (including immediate early genes *Arc* and *Junb*), and 1 DEG (*AC149090*.*1*) in Lamp5 and Vip neurons (**Figure 2a**). In contrast, *Mecp2e1*^*-/+*^ female from experiment 2 single cortical cells showed the strongest transcriptional dysregulation, for a total of 1215 DEGs at P30 (Limma-VoomCC). Interestingly, *Mecp2e1*^*-/+*^ female Pvalb DEGs at P30 had a significant (*p-value* ≤ 0.00075) enrichment of imprinted genes, including *Meg3, Xist, Gnas, Kcnq1ot1, Np1l5, Ntm, Peg3 and Snrpn* (**Figure 2b**), a result that was not observed in males.

**Figure 2.**
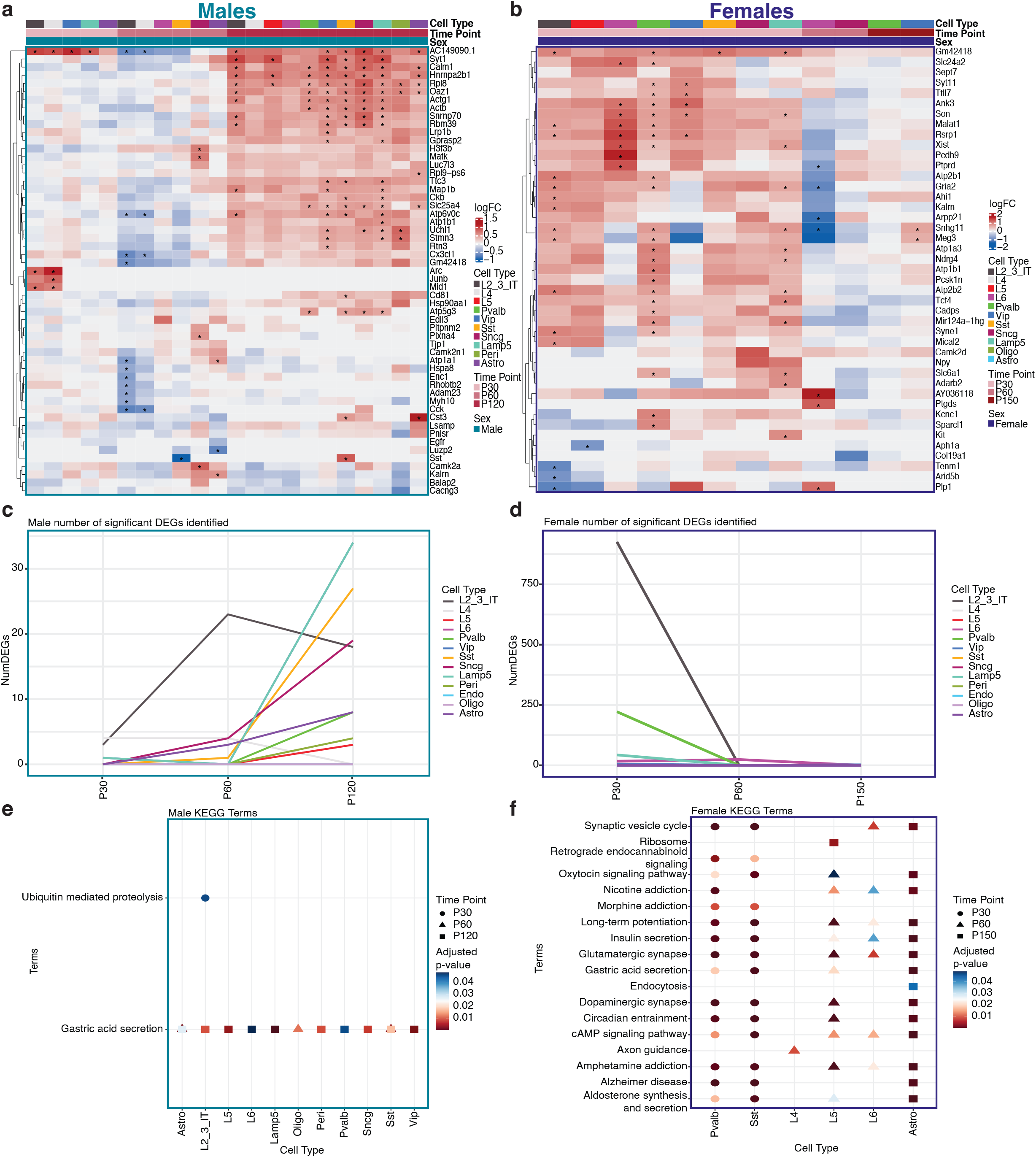
Sexually-dimorphic dynamic patterns of DEGs and KEGG pathway terms across time and cell type. **a**. Heatmap of top 5 differentially expressed genes (DEGs) based on the lowest adjusted p-value ≤ 0.05 comparing male *Mecp2e1*^*-/y*^ and *Mecp2e1*^*+/y*^ cortical cells across timepoints (experiment 1). **b**. Heatmap of top 5 DEGs comparing female *Mecp2e1*^*-/+*^ and *Mecp2e1*^*+/+*^ cortical cells across timepoints (experiment 2). *indicates adjusted p-value ≤ 0.05 (corrected via Benjamini and Hochberg method). **c, d**. Number of DEGs over time at adjusted p-value ≤ 0.05 for experiments 1 and 2, respectively. **e, f**. Dot plots showing the KEGG pathway terms for DEGs (adjusted p-value ≤ 0.1) from each cell type, selected for terms that are persistent over time for experiments 1 and 2, respectively.

At the disease onset P60 timepoint, 73 DEGs were identified in *Mecp2e1*^*-/y*^ males, predominated by 56 DEGs in L2/3 neurons, but also including 7 DEGs in astrocytes, 4 DEGs in L4 and 6 in Scng neurons and 1 DEG in L6 and Sst neurons (**Figure 2c**). *Mecp2e1*^*-/+*^ female cortical cells had 47 DEGs, with 46 in L6 excitatory neurons and 1 DEG in Sncg inhibitory neurons (**Figure 2d**). Further, *Mecp2e1*^*-/+*^ female DEGs at P60 included *AY036118* (+3.47-fold change), *Ptprd, Edil3, Ptgds, Plp1, Atp6v0b, Kcn11ot1, Gria2, Nrxn1, Arpp21, Snhg11. Mecp2e1*^*-/+*^ females had 3 DEGs with 2 in VIP inhibitory neurons and 1 DEG in Pvalb inhibitory neurons.

By the late disease P150 time point, only VIP interneurons contained DEGs in *Mecp2e1*^*-/+*^ cortical cell types, including long non-coding RNAs *Snhg11* (p value = 0.0043) and *Meg3* (p value = 0.017). Remarkably, *Mecp2e1*^*-/+*^ female cortical cells were most transcriptionally dysregulated prior to the onset of symptoms, as the number of DEGs decreased in number as disease symptoms progressed (**Figure 2d**). Overall, *Mecp2e1*^*-/y*^ male DEGs increased in number with disease progression, but *Mecp2e1*^*-/y*^ male cortical cell types had only 199 DEGs across all three time points, which was only 16.3% of the total *Mecp2e1*^*-/+*^ female DEGs (**Figure 2c, d**).

To identify enriched functional pathways connecting RTT transcriptional progression, Kyoto Encyclopedia of Genes and Genomes (KEGG) analysis was performed using DEGs (Limma-VoomCC, p-value ≤ 0.05) from each each cell type. KEGG pathways that were persistent over P30, P60, and P120 in *Mecp2e1*^-/y^ male cortical cells or P30, P60 and P150 in *Mecp2e1*^*-/+*^ female cortical cells are shown (**Figure 2e-f, Supplemental Table 2**). Distinctly different pathway dysregulation was observed between *Mecp2e1*^*-/+*^ females and *Mecp2e1*^-/y^ male cortical cells by two key metrics. First, *Mecp2e1*^-/+^ cortical cell DEGs were enriched for 18 different pathways consistently across time points, compared to only two in *Mecp2e1*^-/y^ males, of which only gastric acid secretion overlaps with *Mecp2e1*^-/+^ pathways. Second, specifically in pre-symptomatic P30 *Mecp2e1*^-/+^ females, Pvalb and Sst neurons shared 14 enriched pathways including synaptic vesicle cycle, retrograde endocannabinoid signaling, oxytocin signaling, morphine addiction, long-term potentiation, insulin secretion, glutamatergic synapse, gastric acid secretion, dopaminergic synapse, circadian entrainment, cAMP signaling pathway, amphetamine addiction, Alzheimer’s disease, and aldosterone synthesis and secretion (**Figure 2f**) compared to only the gastric acid secretion pathway at the disease onset P60 and late disease P120 time point in *Mecp2e1*^-/y^ males (**Figure 2e**). Interestingly, by symptom onset at P60 in *Mecp2e1*^-/+^ females, 6 dysregulated KEGG pathways including nicotine addiction, long-term potentiation, insulin secretion, glutamatergic synapse, cAMP signaling, and amphetamine addiction (found in P30 Pvalb and Sst neurons) were distinctly significantly enriched in L5 and L6 excitatory neurons (**Figure 2f**). At the late disease P150 timepoint, *Mecp2e1*^-/+^ female cortical astrocytes remarkably and distinctly were significantly enriched for 15 out of the 18 total convergent KEGG pathways (**Figure 2f**). While some of the reduced KEGG pathway enrichment in *Mecp2e1* deficient males compared to females could be due to fewer DEGs observed overall and especially at the pre-symptomatic stage in *Mecp2e1*^-/y^ cortical cells, the significant enrichment of ubiquitin mediated proteolysis specifically at *Mecp2e1*^-/y^ P30 when DEGs were fewest (**Figure 2e**) suggests that DEG number is less important than the specificity of gene pathways dysregulated in the female mouse model. We also performed an enrichment analysis for DEGs based on gene length, but did not find evidence to support the previously reported repression of long genes in *Mecp2* deficient neurons^16^ of either sex (**Supplemental Figure 6**).

### Co-expression networks of dysregulated genes within cortical cell types correlate with *Mecp2e1* genotype, time point, sex, body weight, and disease score

To complement the DEG analysis, we performed a systems-biology based approach, High-Definition Weighted Gene Co-expression Network Analysis (hdWGCNA) which is specifically designed for analysis of high dimensional data such as snRNA-seq^17,18^ (**Figure 3**). hdWGCNA groups genes that are co-expressed together into colored modules based on scale-free topology^17–19^ and was used to define nine distinct modules based on co-expression within a network built from transcriptomes of all detected genes from all cell types and experimental conditions. Genes in each module were compared to cell type marker genes to identify modules that uniquely correlate with phenotype (**Supplemental Figure 7**). In co-expression network analysis, we focus on the hub genes, those which are highly connected within each module. Therefore, we determine the eigengene-based connectivity, also known as kME, of each gene. The top 10 ranked co-expressed hub genes were identified per module (**Figure 3a**) and expression enrichment for each cortical cell type was determined, which was distinct from cell type markers (**Figure 3b**). The blue hdWGCNA module corresponded to genes enriched in oligodendrocytes, while the magenta module genes were enriched in L5 and L6 neurons, with *Sez6* and *Nrp1* as hub genes. The brown module included *Grin2a, Grin2b*, and *Camk2a* and the black module included *Slit3* and *Gabrb3* enriched in excitatory neurons (L2-3, L4, L5, L6) showed similar cellular patterns of expression. The green module genes like *Grik1* and *Adarb2* were most highly expressed in inhibitory neurons. In contrast, the turquoise, pink, red and yellow modules are more cell type independent, being enriched in all neuronal subtypes (**Figure 3b**).

**Figure 3.**
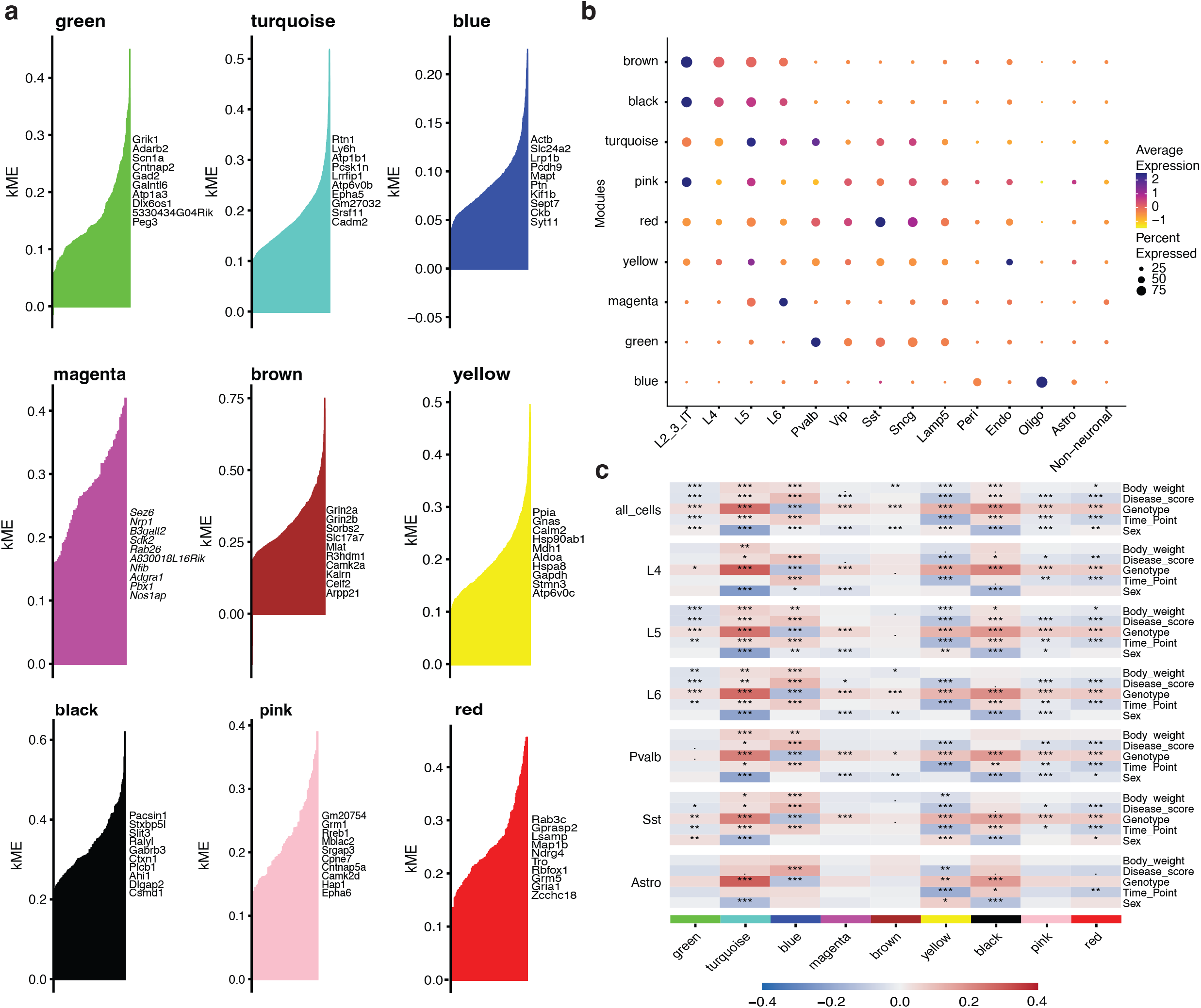
hdWGCNA identifies co-expression networks for each cell type in the mouse cortex that correlated with *Mecp2e1* genotype, disease phenotypes, and sex. **a**. Top 10 hub genes identified for each of the 9 modules generated by hdWGCNA on entire snRNAseq dataset, identified by color. The x-axis are all the genes in each module and the y-axis is the corresponding kME value. **b**. Dot plot of the average gene expression of the top 10 hub genes in each module generated for each cortical cell type. **c**. A heat map of correlations between experimental phenotypes and variables (body weight, disease score, genotype, time point, sex) and averaged gene expression (eigenmode value) for each cell type (cell types not shown are in **Supplemental Figure 7a**). *, **, *** indicates FDR-corrected p-value ≤ 0.05, 0.01, and 0.001, respectively. The color bar shows the Pearson correlation coefficient.

To explore the relationship between cortical co-expression gene networks and disease progression in Rett syndrome, the eigengene value of each sample within each module was correlated with the body weight, disease score, genotype, sex, and disease time point of each mouse. Eigengene values were calculated for all cortical cells, as well as each cell type individually, so that correlations with each variable of interest could be examined for each cell type (**Figure 3c, Supplemental Figure 7)**. While the genes within each module partially overlapped with those that served as cell type markers, the genes within modules were independent from those that defined cell type specificity (**Supplemental Figure 7**). For all cell types, 6 out of the 9 gene modules significantly correlated with all phenotypes and experimental variables, and all modules showed at least one significant correlation (**Figure 3c, top row**). Yet, certain gene set modules such as green correlated with phenotypes in L4, L6 and Sst neurons while the turquoise and blue modules correlated most strongly with phenotypes in all neuronal subtypes (**Figure 3c**). Interestingly, magenta module genes only correlated with genotype in neurons. Astrocytes were distinct in that only blue and yellow modules correlated with both disease score and genotype. While most module-genotype correlations were positive (red), meaning that co-expressed genes in these modules were upregulated in mutant animals, the blue module uniquely was inversely correlated (blue), representing downregulated genes. Interestingly, module-sex associations were frequent but sometimes were absent in specific cell types or time points with strong module-genotype correlations, such as the blue module in L6. Pvalb, and Sst neurons (**Figure 3c**). Further, genes from each module were tested for KEGG pathway enrichment to identify phenotype correlated dysregulated pathways. Many RTT disease progression relevant pathways such as glutamatergic synapse, gabaergic synapse, circadian rhythm and axon guidance were identified (**Supplemental Figure 8**). KEGG analysis for the turquoise module showed enrichment in neurological pathways such as Alzheimer disease and metabolic pathways such as choline metabolism (**Supplemental Figure 8**).

### X chromosome expression mosiacism in female cortical cell populations reveals dynamic non-cell-autonomous transcriptional homeostasis

To examine non-cell-autonomous effects, we considered all *Mecp2* expressing cells within *Mecp2e1*^-/+^ mosaic female cortical cells. Within *Mecp2e1*^-/+^ female snRNA-seq data, we identified 1,146 *Mecp2* expressing cells, of which 539 could be genotyped as WT-expressing and 607 were expressing the *Mecp2e1* mutation (**Supplemental Figure 9**). These cells were clustered based on expression and thus twelve different cell types were identified (**Figure 4a**). To reduce the impact of lower cell counts on DEG calling following parsing, we further grouped the *Mecp2* expressing cells into two broad categories of GABAergic neurons and glutamatergic neurons (**Figure 4b**). A summary of the number of DEGs in each of these three broader cell type categories (glutamatergic, GABAergic, non-neuronal for each of the five experimental comparisons (**Fig 1d**) is shown in **Table 1**. LimmaVoom was used for DEG calling of experiments 3, 4 and 5. As expected based on random XCI, both cell populations (Mecp2_MUT and Mecp2_WT), were randomly represented in all cell types (**Figure 4a, b, c**). Cells from the males were also parsed, showing 184 Mecp2_MUT in the *Mecp2e1*^-/y^ and 175 Mecp2_WT in the *Mecp2e1*^+/y^ (**Supplemental Figure 9**).

**Table 1.** Summary of LimmaVoom DEG numbers resulting from the parsing of mutant- and WT-expressing cell type categories in all comparison experiments.

| Exp # | Cell type | #UP DEGs | #DOWN DEGs | Time point | Animal (n) | Total sig DEGs | Notes |
| --- | --- | --- | --- | --- | --- | --- | --- |
| 1 | Glutamatergic | 7 | 0 | P30 | 4 | 7 | Comparing all cortical glutamatergic cells between mutant male *Mecp2e1-/y* to WT male *Mecp2e1+/y* |
| 1 | GABAergic | 2 | 0 | P30 | 4 | 2 | Comparing all cortical GABAergic cells between mutant male *Mecp2e1-/y* to WT male *Mecp2e1+/y* |
| 1 | Non-neuronal | 0 | 0 | P30 | 4 | 0 | Comparing all cortical non-neuronal cells between mutant male *Mecp2e1-/y* to WT male *Mecp2e1+/y* |
| 1 | Glutamatergic | 0 | 27 | P60 | 4 | 27 | Comparing all cortical glutamatergic cells between mutant male *Mecp2e1-/y* to WT male *Mecp2e1+/y* |
| 1 | GABAergic | 4 | 1 | P60 | 4 | 5 | Comparing all cortical GABAergic cells between mutant male *Mecp2e1-/y* to WT male *Mecp2e1+/y* |
| 1 | Non-neuronal | 2 | 1 | P60 | 4 | 3 | Comparing all cortical non-neuronal cells between mutant male *Mecp2e1-/y* to WT male *Mecp2e1+/y* |
| 1 | Glutamatergic | 21 | 0 | P120 | 4 | 21 | Comparing all cortical glutamatergic cells between mutant male *Mecp2e1-/y* to WT male *Mecp2e1+/y* |
| 1 | GABAergic | 121 | 1 | P120 | 4 | 126 | Comparing all cortical GABAergic cells between mutant male *Mecp2e1-/y* to WT male *Mecp2e1+/y* |
| 1 | Non-neuronal | 8 | 0 | P120 | 4 | 8 | Comparing all cortical non-neuronal cells between mutant male *Mecp2e1-/y* to WT male *Mecp2e1+/y* |
| 1 | **Total** | **165** | **30** |  | **32** | **195** |  |
| 2 | Glutamatergic | 31 | 913 | P30 | 4 | 944 | Comparing all cortical glutamatergic cells between mutant female *Mecp2e1-/+* to WT female *Mecp2e1+/+* |
| 2 | GABAergic | 236 | 35 | P30 | 4 | 271 | Comparing all cortical GABAergic cells between mutant female *Mecp2e1-/+* to WT female *Mecp2e1+/+* |
| 2 | Non-neuronal | 0 | 0 | P30 | 4 | 0 | Comparing all cortical non-neuronal cells between mutant female *Mecp2e1-/+* to WT female *Mecp2e1+/+* |
| 2 | Glutamatergic | 13 | 11 | P60 | 4 | 24 | Comparing all cortical glutamatergic cells between mutant female *Mecp2e1-/+* to WT female *Mecp2e1+/+* |
| 2 | GABAergic | 0 | 0 | P60 | 4 | 0 | Comparing all cortical GABAergic cells between mutant female *Mecp2e1-/+* to WT female *Mecp2e1+/+* |
| 2 | Non-neuronal | 0 | 0 | P60 | 4 | 0 | Comparing all cortical non-neuronal cells between mutant female *Mecp2e1-/+* to WT female *Mecp2e1+/+* |
| 2 | Glutamatergic | 0 | 0 | P150 | 8 | 0 | Comparing all cortical glutamatergic cells between mutant female *Mecp2e1-/+* to WT female *Mecp2e1+/+* |
| 2 | GABAergic | 2 | 0 | P150 | 8 | 2 | Comparing all cortical GABAergic cells between mutant female *Mecp2e1-/+* to WT female *Mecp2e1+/+* |
| 2 | Non-neuronal | 0 | 0 | P150 | 8 | 0 | Comparing all cortical non-neuronal cells between mutant female *Mecp2e1-/+* to WT female *Mecp2e1+/+* |
| 2 | **Total** | **282** | **959** |  | **32** | **1241** |  |
| 3 | Glutamatergic | 17 | 782 | P30 | 4 | 799 | Comparing WT-expressing glutamatergic cells from mutant female *Mecp2e1-/+* to WT-expressing cells from WT female *Mecp2e1+/+* |
| 3 | GABAergic | 2 | 393 | P30 | 4 | 395 | Comparing WT-expressing GABAergic cells from mutant female *Mecp2e1-/+* to WT-expressing cells from WT female *Mecp2e1+/+* |
| 3 | Glutamatergic | 835 | 5 | P60 | 4 | 840 | Comparing WT-expressing glutamatergic cells from mutant female *Mecp2e1-/+* to WT-expressing cells from WT female *Mecp2e1+/+* |
| 3 | GABAergic | 5 | 0 | P60 | 4 | 5 | Comparing WT-expressing GABAergic cells from mutant female *Mecp2e1-/+* to WT-expressing cells from WT female *Mecp2e1+/+* |
| 3 | Glutamatergic | 2 | 41 | P150 | 8 | 43 | Comparing WT-expressing glutamatergic cells from mutant female *Mecp2e1-/+* to WT-expressing cells from WT female *Mecp2e1+/+* |
| 3 | GABAergic | 1 | 43 | P150 | 8 | 44 | Comparing WT-expressing GABAergic cells from mutant female *Mecp2e1*<sup>-/+</sup> to WT-expressing cells from WT female *Mecp2e1*<sup>+/+</sup> |
| **3** | **Total** | **862** | **1264** |  | **32** | **2126** |  |
| 4 | Glutamatergic | 3 | 574 | P30 | 4 | 577 | Comparing mutant-expressing glutamatergic cells from female *Mecp2e1*<sup>-/+</sup> to WT-expressing cells from female WT *Mecp2e1*<sup>+/+</sup> |
| 4 | GABAergic | 9 | 984 | P30 | 4 | 993 | Comparing mutant-expressing GABAergic cells from female *Mecp2e1*<sup>-/+</sup> to WT-expressing cells from female WT *Mecp2e1*<sup>+/+</sup> |
| 4 | Glutamatergic | 1251 | 8 | P60 | 4 | 1259 | Comparing mutant-expressing glutamatergic cells from female *Mecp2e1*<sup>-/+</sup> to WT-expressing cells from female WT *Mecp2e1*<sup>+/+</sup> |
| 4 | GABAergic | 0 | 0 | P60 | 4 | 0 | Comparing mutant-expressing GABAergic cells from female *Mecp2e1*<sup>-/+</sup> to WT-expressing cells from female WT *Mecp2e1*<sup>+/+</sup> |
| 4 | Glutamatergic | 4 | 8 | P150 | 8 | 12 | Comparing mutant-expressing glutamatergic cells from female *Mecp2e1*<sup>-/+</sup> to WT-expressing cells from female WT *Mecp2e1*<sup>+/+</sup> |
| 4 | GABAergic | 0 | 4 | P150 | 8 | 4 | Comparing mutant-expressing GABAergic cells from female *Mecp2e1*<sup>-/+</sup> to WT-expressing cells from female WT *Mecp2e1*<sup>+/+</sup> |
| **4** | **Total** | **1267** | **1578** | **0** | **32** | **2845** |  |
| 5 | Glutamatergic | 0 | 0 | P30 | 2 | 0 | Comparing mutant-expressing glutamatergic cells from female *Mecp2e1*<sup>-/+</sup> to WT-expressing cells from female *Mecp2e1*<sup>-/+</sup> |
| 5 | GABAergic | 0 | 0 | P30 | 2 | 0 | Comparing mutant-expressing GABAergic cells from female *Mecp2e1*<sup>-/+</sup> to WT-expressing cells from female *Mecp2e1*<sup>-/+</sup> |
| 5 | Glutamatergic | 3 | 0 | P60 | 2 | 3 | Comparing mutant-expressing glutamatergic cells from female *Mecp2e1*<sup>-/+</sup> to WT-expressing cells from female *Mecp2e1*<sup>-/+</sup> |
| 5 | GABAergic | 0 | 0 | P60 | 2 | 0 | Comparing mutant-expressing GABAergic cells from female *Mecp2e1*<sup>-/+</sup> to WT-expressing cells from female *Mecp2e1*<sup>-/+</sup> |
| 5 | Glutamatergic | 6 | 0 | P150 | 4 | 6 | Comparing mutant-expressing glutamatergic cells from female *Mecp2e1*<sup>-/+</sup> to WT-expressing cells from female *Mecp2e1*<sup>-/+</sup> |
| 5 | GABAergic | 1 | 0 | P150 | 4 | 1 | Comparing mutant-expressing GABAergic cells from female *Mecp2e1*<sup>-/+</sup> to WT-expressing cells from female *Mecp2e1*<sup>-/+</sup> |
| **5** | **Total** | **10** | **0** | **0** | **16** | **10** |  |

**Figure 4.**
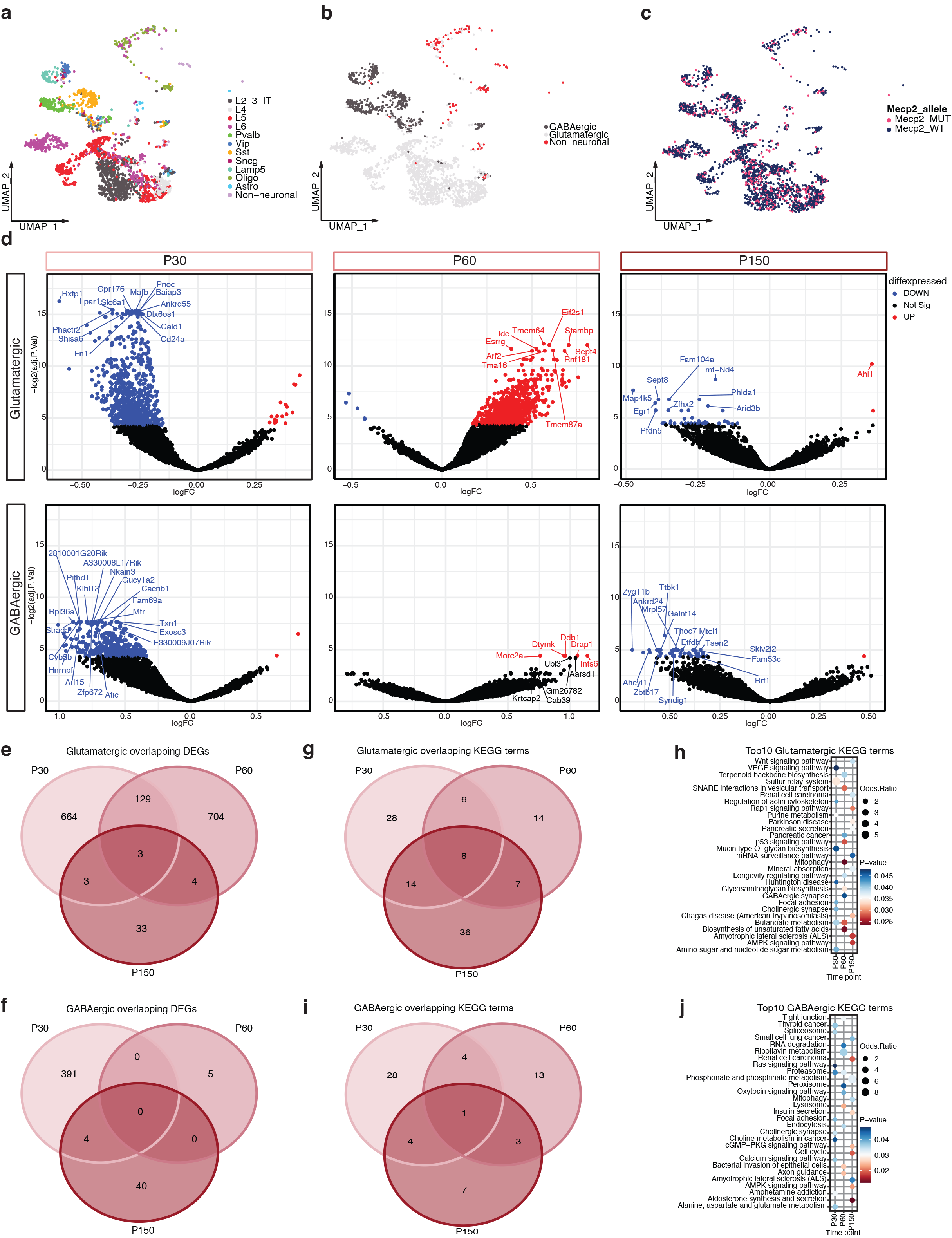
Dynamic non-cell-autonomous effects on differentially expressed genes and KEGG pathways over disease progression. As shown in Experiment #3 (Fig. 1d), we compared WT cells from *Mecp2e1*^*-/+*^ female with WT cells from *Mecp2e1*^*+/+*^ glutamatergic and GABAergic neurons longitudinally. **a**. UMAP plot of cell types indentified in the mosaic females **b**. UMAP plot of the female cortices showing the clustering of the broad cell type categories. **c**. UMAP plot of mosaic female cells parsed by *Mecp2* allele **d**. Volcano plots showing differentially expressed genes (DEGs) of the mouse cortical neurons contrasting WT cells from WT *Mecp2e1*^*+/+*^ females and WT cells from *Mecp2e1*^*-/+*^ mosaic females. **e**,**f**. Venn diagrams of overlapping glutamatergic and GABAergic DEGs respectfully over time. **g**,**i**. Venn diagrams of significant KEGG terms of glutamatergic and GABAergic neurons over time. **H**. Top 10 KEGG terms of glutamatergic neurons over time. **j** Top 10 KEGG terms of GABAergic neurons over time.

In order to examine non-cell-autonomous effects over the disease progression in a cell category specific manner, we followed the third experiment design (**Figure 1d**) and compared the WT expressing cells from the *Mecp2e1*^-/+^ mosaic females to the WT expressing cells from the *Mecp2e1*^+/+^ females (**Figure 4d, Table 1**). At P30, both glutamatergic and GABAergic WT-expressing cells from *Mecp2e1*^*-/+*^ showed a large number of significant downregulated genes (blue) but a low number of upregulated genes (red), despite these cell populations being WT-expressing. These differences in gene expression were likely due to non-cell-autonomous effects of the *Mecp2e1* mutation on nearby WT-expressing cells. Further evidence was obtained from the experimental comparison from experiment 5, where mutant-expressing glutamatergic and GABAergic neurons from female *Mecp2e1*^*-/+*^ were compared to WT-expressing cells from female *Mecp2e1*^*-/+*^, resulting in only 10 DEGs, compared to 862 in experiment 3 (non-cell-autonomous WT vs WT) (**Table 1**). This non-cell-autonomous effect was dynamic over time, as glutamatergic cells showed mostly upregulated genes with only a few downregulated genes at P60, while GABAergic cells only showed upregulated genes (**Figure 4d**). Interestingly, at the late disease stage P150, the number of dysregulated genes were diminished and primarily back to being downregulated, indicating a dynamic process of non-cell-autonomous effects across disease progression. In order to test this hypothesis, we overlapped the significant (LimmaVoom adjusted p-value ≤ 0.05) DEGs from each cell type and time point (**Figure 4e, 4f**). In glutamatergic cells, the largest overlap (129 DEGs) was between P60 and P150 and only 3 DEGs in common to all three time points (**Figure 4e**). Similar results were seen in GABAergic neurons, where DEGs were predominently unique to each time point (**Figure 4f**).

In order to look for functional pathway enrichments of non-cell-autonomous effects of *Mecp2e1*^-/+^ mosaicism, KEGG analysis on significant DEGs (p-value ≤ 0.05) from glutamatergic and GABAergic neurons was performed and significant (adjusted p-value ≤ 0.05) terms overlapped across time (**Figure 4g, 4i**). Non-cell-autonomous DEGs from glutamatergic cells were enriched for 8 terms that were shared across all disease stages which include Parkinson, Alzheimer, and Huntington diseases, as well as homeostatic pathways of retrograde endocannabinoid signaling, ubiquitin mediated proteolysis, oxidative phosphorylation, and protein processing in endoplasmic reticulum, while the P60 time point was uniquely enriched for terms such as GABAergic synapse and SNARE interactions in vesicular transport (**Figure 4h**). Further, glutamatergic cells showed molecular dysregulation associated with MeCP2 activity such as mRNA surveillance pathway, cholinergic synapse, and AMPK signaling pathway (**Figure 4h**). In contrast, GABAergic cells shared axon guidance as an enriched pathway common across all time points (**Figure 4i**). Other interesting RTT related pathways included metabolism and energy related terms such as riboflavin metabolism, phosphonate and phosphinate metabolism, choline metabolism, and alanine, aspartate, and glutamate metabolism (**Figure 4j**).

In order to compare these non-cell-autonomous effects to cell-autonomous effects over the disease progression in a cell category specific manner, we followed the fourth experiment design (**Figure 1d**) and compared the MUT *Mecp2* expressing cells from the *Mecp2e1*^-/+^ mosaic females to the WT *Mecp2* expressing cells from the *Mecp2e1*^+/+^ females (**Supplemental Figure 10a**). Similar to the results of experiment 3, glutamatergic and GABAergic significant DEGs were predominantly time point specific (**Supplemental Figure 10b-c**). Glutamatergic cells had 12 significant KEGG pathways shared over time while GABAergic cells had 6 significant terms both containing RTT related pathways such as mRNA surveillance and circadian rhythm (**Supplemental Figure 10d-g**). In order to examine if the dysregulated KEGG pathways are shared between experiment 3 and experiment 4, a comprehensive overlap test was performed showing majority of the pathways are unique to each experiment and each time, with the glutamatergic P150 KEGG pathways from non-cell-autonomous DEGs outnumbering those of cell-autonomous (17 in exp 3 vs 1 in exp 4) (**Supplemental Figure 11**).

Lastly, we examined non-cell-autonomous effects by comparing MUT-expressing to WT-expressing cells within the mosaic *Mecp2e1*^-/+^ females, as described in experiment 5 (**Figure 1d**). Overall, glutamatergic and GABAergic had only a few genes dysregulated which were mostly at P150 when analyzed separately (**Supplemental Figure 12a**). For higher statistical power in KEGG term enrichment, DEGs glutamateric and GABAergic cells were each combined across time points, revealing dysregulated retrograde endocannabinoid signaling and other pathways (**Supplemental Figure 12b-c**). The top10 enriched KEGG pathways when both glutamatergic and GABAergic cells were combined across all time points included pathways involved in cell signaling and addiction (**Supplemental Figure 12d**). The differences between WT-expressing and MUT-expressing cells within mosaic females in experiment 5 were far less than the differences between WT-expressing cells in mosaic *Mecp2e1*^*-/+*^ females compared to WT cells in *Mecp2e1*^*+/+*^ females in experiment 4. Together, these analyses demonstrate that transcriptional dysregulation across disease progression in mosaic *Mecp2e1*^-/+^ females is dynamic and predominated by non-cell-autonomous effects on homeostatic gene pathways.

### Human RTT cortical cell transcriptional dysregulation is recapitulated by the female but not the male RTT mouse model

To examine how closely *Mecp2e1*^-/+^ mice phenocopy Rett syndrome (RTT) at the cellular transcriptome level, we examined the relationship between altered transcript levels by cell type in *Mecp2e1* deficient and human *MECP2*^*-/+*^ cortices. Thus sn-RNA seq analysis was performed on eight *MECP2*^*-/+*^ (RTT) and eight age matched control female cortex samples from post-mortem human brains (**Figure 5a**). Nine neuronal and six non-neuronal cell type clusters could be assigned from these human cortices based on 3,000 gene markers from the Bakken Trygve et al. dataset^20^ (**Figure 5b**). Cell type labeling based on scTransform containing elevated expression of at least three cell marker genes was validated (**Figure 5c**). DEG analysis via limmaVoomCC compared *MECP2*^*-/+*^ to *MECP2*^*+/+*^ cortical cells, resulting in cell type-specific dysregulated genes (**Figure 5d**). Importantly, the top 20 upregulated DEGs identified by LimmaVoomCC at the adjusted *p*-value ≤ 0.05 level in female *MECP2*^*-/+*^ cortical cells were also significant DEGs (adjusted *p* value ≤ 0.05) in *Mecp2e1*^*-/+*^ female mouse cortices with 14 gene transcripts out of 20 upregulated (**Figure 5e**). Similarly, of the top 20 Rett female mouse cortical LimmaVoomCC DEGs that were significantly downregulated (adjusted *p* value ≤ 0.05), the homologous *Mecp2e1*^*-/+*^ female gene transcripts were also downregulated (**Figure 5e**). In contrast, there were very few overlapping DEGs between human RTT and *Mecp2e1*^*-/y*^ male cortical cell transcriptomes (**Figure 5f**). This demonstrates that *Mecp2e1*^*-/+*^ female mice are a better model for the dynamic transcriptomic dysregulation due to cellular complexities in Rett syndrome disease progression.

**Figure 5.**
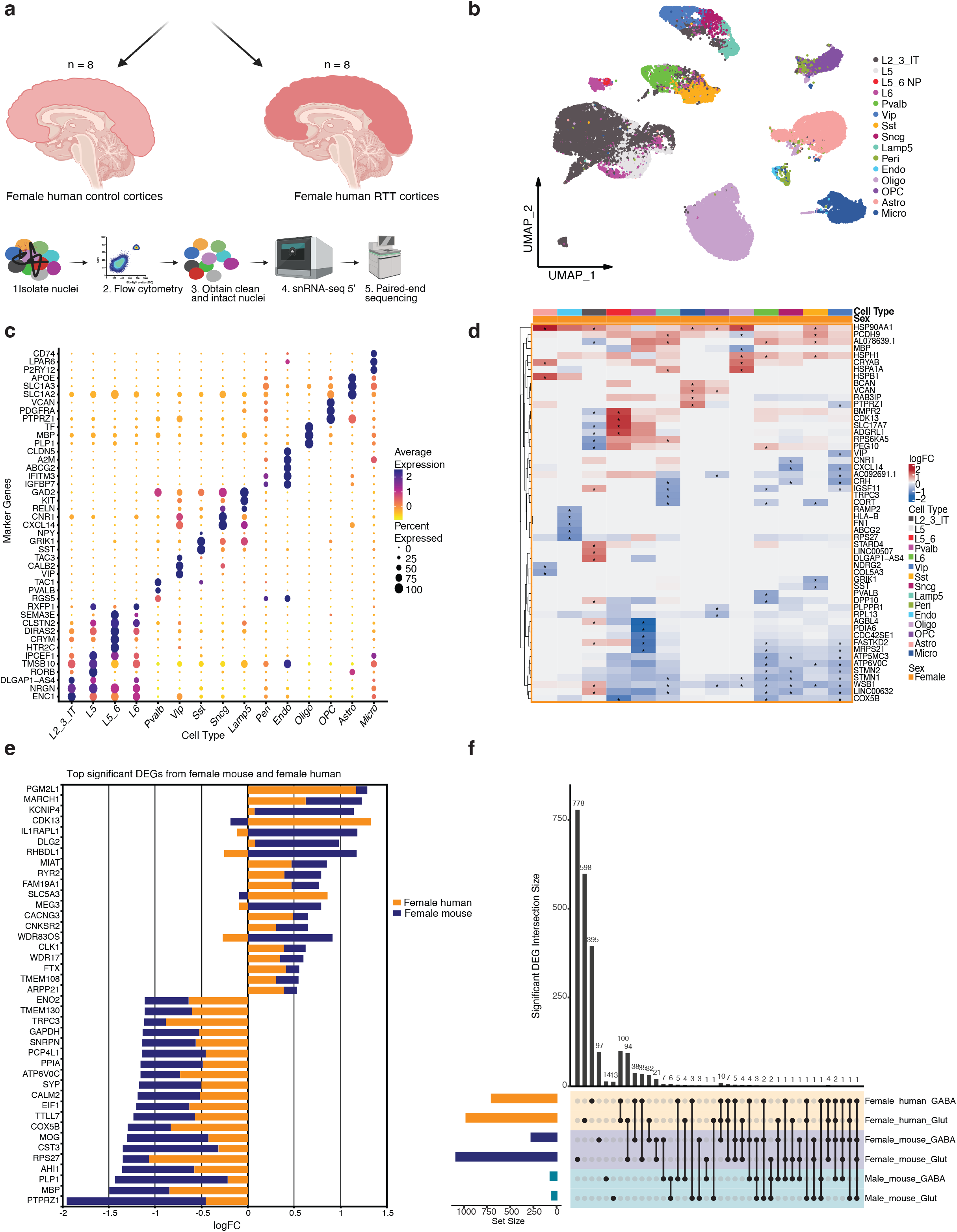
Human RTT cortical neurons share transcriptional dysregulation specifically with *Mecp2e1*^*-/+*^ mosaic female mice. **a**. A schematic of postmortem human RTT cortices and age/sex matched control cortices. **b**. UMAP of the unsupervised clustering of cell types identified in the human cortices (n = 39336 cells post QC). Cell type labels were transferred from Bakken Trygve et al. 2021^20^. **c**. Top gene markers for each cell type in the human cortex. **d**. Heatmap of top differentially expressed genes (DEGs) for human female cortices. *indicates adjusted p-value ≤ 0.05. **e**. Bar graph showing overlapping of the top significant upregulated and downregulated genes by logFC in female mouse and female human. **f** Upset plot showing overlap of the significant DEGs from both GABAergic and glutamatergic neurons in female human, female mouse, and male mouse.

## Discussion

This study advances our understanding of RTT, offering insights into sex-specific, cell type-dependent, and disease stage-associated transcriptional dysregulation resulting from the cellular complexities related to the X-linked dominant inheritance of *MECP2/Mecp2* mutation. This longitudinal analysis of single cortical cell transcriptomes during the gradual progression of disease symptoms in the *Mecp2e1*^*-/+*^ mouse model of RTT provided several new findings critical to understanding and treatment of human RTT. First, we demonstrated that the female *Mecp2e1*^*-/+*^ mice are inherently different, not simply less severe, in their transcriptional dysregulation compared to mutant *Mecp2*^*-/y*^ males that completely lack *Mecp2e1*. Second, we identified transcriptionally dysregulated gene pathways across cell types in female *Mecp2e1*^*-/+*^ cortices that were significantly associated with progression of multiple disease phenotypes over time. Third, we showed that non-cell-autonomous effects in mosaic female *Mecp2e1*^*-/+*^ mice are responsible for the homeostatic gene pathway dysregulations observed dynamically over time. Lastly, and most important for translational relevance, we showed that female mosaic *Mecp2e1* mutant mice better recapitulate the transcriptional dysregulation observed in human RTT cortical cells than *Mecp2* null males, and may help explain the complexities of progressive and regressive stages of disease in RTT girls.

The earliest studies examining the effect of MeCP2 levels on transcription in brain relied on bulk RNA-seq comparing male *Mecp2* null to wild-type controls yielded few differentially expressed gene (DEG) transcripts^21–24^. However, bulk analyses of transcript levels in *Mecp2* null compared to *Mecp2* duplication mouse brain revealed 2582 altered DEGs in hypothalamus^25^, 1180 DEG transcripts in cerebellum^26^, and 1060 DEGs in amygdala^27^. Interestingly, analysis of transcripts in individual brain cell types yielded non-overlapping lists of DEGs suggesting that bulk tissue DEG analysis suffers from a “dilution effect” potentially masking DEGs^28^. While these initial studies comparing *Mecp2* null to wild-type and *Mecp2* duplication control male brains can reveal gene targets of MeCP2 *in vivo, Mecp2*^*-/+*^ female mice are the relevant model for understanding RTT, where brain cell autonomous and non-autonomous effects require analysis of individual cells and cell types.

For autosomal genes, heterozygous mutations are expected to show reduced phenotypic severity than the homozygous state, but for X-linked genes, there is the added complication of random XCI that creates epigenetic mosaicism within cell populations. We were able to utilize sn-RNA seq 5’ in the *Mecp2e1* mouse model to parse by both cortical cell type and mutation to improve understanding of transcriptional dysregulation in RTT. Our results can both help confirm certain aspects of previous bulk transcriptomic studies and help explain some of the prior discrepancies between bulk transcriptomic studies in RTT mouse models. A study using bulk RNA-seq on 7 week-old *Mecp2* null mice showed 48 genes upregulated and 32 genes down-regulated in pathways such as circadian entrainment that are consistent with our single nucleus data, despite the lower overall number of DEGs identified^29^. We identified circadian entrainment as a homeostatic gene pathway dynamically dysregulated in Pvalb and Sst at P30, L5 excitatory neurons at P60, and astrocytes at P150. To date, there has been one prior study conducted using single nucleus RNA-seq in both RTT brain and a mouse model^16^. Renthal *et al* used *Mecp2* null cortex at a single time point (8 weeks for males, 12-20 weeks for females) and compared to human RTT cortex with *MECP2* 255X^16^. Our studies were consistent in finding both up- and down-regulated genes with MeCP2 deficiency across cell types and in finding evidence for non-cell-autonomous gene dysregulation, but inconsistent in demonstrating a significant effect of MeCP2 deficiency on repression of long genes. Differences between the study designs, including genetic mouse model, time points, statistical approaches for DEGs, and single cell technonology (iDrops versus 5’ V2 technology) could explain the discrepancies. We specifically designed the current study to overcome some of the prior technical limitations, including improvement in signal to noise ratio^30^, higher number of genes detected per nucleus, higher UMI per nucleus, and higher number of cells analyzed^31^. Further, we used 3,000 marker genes from the Allen brain atlas cortex single nucleus dataset to label cell types (compared to one marker gene per cell) and used five different statistical approaches to robustly identify differentially expressed genes.

To understand how transcriptional dysregulation in RTT cortex was related to symptom progression, our study uniquely utilized a longitudinal study design and systems biology approaches to correlate networks of dysregulated gene expression patterns with disease phenotypes over time. Remarkably, these disease-relevant gene networks were not specific to individual cell types, but instead were enriched in pathways also dysregulated in neurodegenerative disorders and addiction pathways that regulate brain homeostasis across cell types, including metabolism, circadian entrainment, and retrograde endocannabinoid signaling. Previous studies had shown a link between MeCP2 and addiction^32^ that involve *Arc* and *Junb* transcription consistent with our results in *Mecp2e1*^*-/y*^ cortical cells^33^ and KEGG pathways enriched in *Mecp2e1*^*-*^ _*/+*_ cortical cells. A prior study showing that selective deletion of *Mecp2* from excitatory neurons had no effect on excitatory transmission, but reduced inhibitory synapse numbers and neurotransmission in the somatosensory and prefrontal cortex^34^ is consistent with our results showing a spread of dysregulated gene pathways from excitatory to inhibitory neurons. A more recent study investigating neurons and astrocytes found KEGG pathways such as calcium signaling pathway and Rap1 signaling pathway were enriched in RTT, consistent with our findings^35^.

While non-cell autonomous effects have been previously noted in RTT mouse models^6–8^, our comprehensive analyses of cellular transcriptomes over disease progression implicates these effects as a central and defining feature of transcriptional dysregulation in RTT mosaic females. Sun et al. argue that the abnormal morphologies of neurons and astrocytes in human RTT are caused by non-cell-autonomous effects driven by altered gene expression and enriched energy related KEGG pathways consistent with our findings from experiment 2^29^. Defects in signaling pathways suggests RTT disease progression is not caused exclusively by autonomous transcriptional changes in individual cells, but rather due to a failure of wild-type *MECP2* expressing cells to compensate for mutant *MECP2* expressing cells.

Since RTT in humans almost exclusively affects females, our results have important implications for translational medicine. First, pre-clinical models for testing new therapies should be female and construct-relevant, ideally modeling actual human RTT mutations. While male *Mecp2* null models provide important basic insights into MeCP2 function^5^, we clearly demonstrate that the *Mecp2e1* deficient males do not recapitulate the transcriptional dysregulation observed in RTT human cortical cells as well as their female mutant littermates. Furthermore, the non-cell-autonomous dynamic waves of dysregulation in WT-expressing cortical neurons may help explain why human RTT symptoms appear gradually and are staggered in a series of regressions followed by plateaus. Our results showing that transcriptional dysregulation appears pre-symptomatic in female *Mecp2e1* mutant across multiple cortical cell types suggest that diagnosis and treatment should ideally begin as early as possible, potentially by including *MECP2* mutations in newborn screening panels. To date the only drug in the market for RTT is trofenitide which is based on IGF-1^36^ a growth factor previously used for diseases such as Laron syndrome and liver cirrhosis^37^. The overlap with other neurologic disease pathways including oxidative phosphorylation suggests that some existing drugs for neurodegenerative disorders could potentially be repurposed to counteract some of the RTT non-cell-autonomous transcriptional dysregulations in pathways regulating homeostasis. Conversely, the molecular pathogenesis of RTT may provide insights for understanding epigenetic regulation of transcriptional homeostasis of gene pathways relevant to common neurodegenerative and addiction disorders.

## Methods

### Single nuclei isolation for mouse and human post-mortem cortex

Mecp2-e1 and control mice were sacrificed by carbon dioxide inhalation just prior to brain removal. Cerebral cortex was removed from each brain from the mice. About 10mg of cerebral cortex tissue was isolated from human post-mortem and control samples. Single nuclei were prepared from the left hemisphere cortex according to a previously established protocol Martelotto (https://cdn.10xgenomics.com/image/upload/v1660261285/support-documents/CG000124_Demonstrated_Protocol_Nuclei_isolation_RevF.pdf). Briefly, a 3.0 mm^2^ section of cortex was removed from each mouse brain. Both mouse and human brain tissue were minced with a scalpel then homogenized in 0.5 mls of nuclei lysis buffer with RNAse inhibitor (Roche, Indianapolis, ID) then transferred to a larger tube with an additional 1.0 ml of nuclei lysis buffer, mixed then incubated on ice for 5 minutes. Nuclei were filtered from the lysate using a 70 µM FlowMi cell strainer (Sp-Belart, Wayne, NJ). Nuclei were pelleted at 4°C for 5 minutes at 500xG, resuspended in 1.5 ml of nuclei wash buffer, incubated for 5 minutes. Nuclei were then pelleted again as above then washed twice in nuclei wash and resuspension buffer then filtered with a 35 µM FlowMi filter (Sp-Belart, Wayne, NJ) then resuspended in nuclei wash and resuspension buffer with 5 ugs/ml DAPI and assayed on a Countess cell counter to determine concentration and nuclear integrity (Fisher Scientific, Waltham, MA). Nuclei were then sorted to remove debris and nuclear aggregates on a MoFlow Astrios cell sorter (Beckman-Coulter, Brea, CA). Approximately,150,000 nuclei per sample were sorted and stored on ice prior to sn-RNA seq 5’ library generation.

### Single nuclei-RNA sequencing

Single Cell 5’ Library & Gel Bead Kits (10x Genomics, Pleasanton, CA) were used to prepare cDNA and generate bar coded and indexed snRNA-seq 5’ libraries according to the manufacturers protocol. 10,000 nuclei per sample were targed. snRNA-seq 5’ libraries were balanced using a Kapa library quantification kit (Roche, Indianapolis, IN) and pooled to generate 150 base pair, paired end sequences from using a NovaSeq S4 sequencer (Illumina, San Diego, CA). Mouse cortices had about 75,000 reads per cell on average and 240,437,728 reads per sample on average. Human cortices had about 50,000 reads per cell on average and 300,000,000 reads per sample on average.

### Pre-processing and quality control

Cellranger v.2.0.2 was used to aligned the mouse raw reads to mm10-1.2.0 reference genome and the human raw reads to GRCh38 human reference genome. Cell by gene count matricies were used to create a Seurat object using Seurat_4.3.0.1 in R 4.2.2. Mouse samples were filtered with the critera that cells should have less than 7% mitochondrial, greater than 200 and less than 5,625 genes and greater than 208 and less than 16,300 UMI respectively.

### Cell type identification by dimentionality reduction

The expression counts were log transformed and normalized via Seurat 4.3.0.1. Information about the samples such as sex, genotype, time point, disease score, body weight and *Mecp2e1* expressionallele were all added to the metadata. Single cell mouse and human cortex data from the Allen brain institute were used as a reference for cell type labeling^38,39^ both data sets separately. scTransform was used to align cell types and transfer labels over to the Rett data. Cell marker test was performed for validating the cell type labeling. Dot plots showing validation of the cell type markers were created via scCustomize 2.1.1 (10.5281/zenodo.5706430).

### DEG analysis

A total of five different DEG analysis methods were used to evaluate the best method for comparing mutant samples to WT samples in a cell-type-specific manner. EdgeR, Limma, and DeSeq2 yielded inconsistent DEGs (**Supplemental Figure 1**). For experiments 1 and 2, low expressing genes were filtered out . Low expressing was defined by expression in less than 25% of cells of a given cell type. LimmaVoomCC was used on the remaining high expressing genes to deteremine differentially expressed genes while considering cells from the same mouse will have correlated expression. For the low expressing genes, DEsingle was used for DEG analysis on genes that are not as robustly expressed (expressed in <25% of cells of type). For experiments 3, 4, and 5, LimmaVoom was exclusively used to identify differentially expressed genes. For each of the DEG experiments, the number of cells were normalized by downsampling. Parameters for all DEG analysis are available in the github repository.

### KEGG analysis

DEGs with a p-value of ≤0.05 from each of the experiments were used as the input for KEGG analysis. This was performed using the R package enrichR 3.2. The top 10 KEGG terms were determined based on p-value for all experiments. We also included gene ontology analysis using the same DEGs.

### hdWGCNA analysis

Cells from both males and females in the processed Seurat object were used as the input for hdWGCNA analysis. We also included phenotype data such as disease score. The criteria for the fraction of cells that a gene needs to be expressed in order to be included was set at 5%. The network type used is signed with a softpower of 0.8. A total of 9 modules were produced and scores for each module was computed using UCell method. Standard pipeline for hdWGCNA 0.2.4 were followed and the parameters are available in the github repository.

### WT and mutant cell parsing in mosaic female mouse corticies

All *Mecp2* reads were extracted from the raw fastq files generated from each individual sample. abBLAST 3.0 and BWA 0.7.17 mem were used in conjunction to extract *Mecp2* reads (alleler.py). The reference used for alignment was 100 bp of the *Mecp2* gene; 50 bp upstream of the exon1 start codon and 50 bp downstream. With the aligned reads, the number of mutant (TTG) and wild type (ATG) start codons were counted using alleler.py. Each read also contains the cell barcode and UMI information which was used to add the mutant cell and wild-type cell information back to the Seurat object as metadata.

### Overlap of human and mouse DEGs and KEGG pathways

LimmaVoomCC DEGs from both human cell types and mouse cell types were filtered at adjusted p-value ≤ 0.05. Significant human DEGs were overlapped with female mouse and male mouse respectively. GeneOverlap 1.38.0 package was used to perform a Fisher’s exact test to determine the significance of overlapped genes. The same overlap approach was performed to determine the significant overlapping mouse and human KEGG pathways.

## Data availability

Raw data is in the process of being uploaded to GEO

### Code availablility

The analysis pipeline for the study is available at: https://github.com/osmansharifi/snRNA-seq-pipeline **(currently set to private)**

## Acknowledgements

We would like to thank Bridget McLaughlin, for her expertise in flow cytometry analysis and cell sorting, Lutz Froenicke, and Diana Burkart-Waco, for their expertise with 10X genomics sn-RNA seq protocols.

## Funding

NIH NIAA grant 1R01AA027075 to Janine M. LaSalle, and NIH Shared Instrumentation Grant 1S10OD010786-01 to the UC Davis DNA technologies core. This project was supported by the University of California Davis Flow Cytometry Shared Resource Laboratory with funding from the NCI P30 CA093373 (Comprehensive Cancer Center), and S10 OD018223 (Astrios Cell Sorter) grants, with technical assistance from Bridget McLaughlin and Jonathan Van Dyke. This has been made possible in part by grants from the National Institute of Child Health and Human Development (P50 HD103526).

**Supplemental Table 1.** Table containing all significant up and down regulated DEGs (LimmaVoomCC) from experiment 1 and 2. Table contains gene name, logFC, adjusted p-value, sex, cell type and time point information.

**Supplemental Table 2.**Table containing all significant KEGG pathways from experiment 1 and 2. Table contains Term, overlap, odds ratio, adjusted p-value, Genes contained in the pathway, sex, cell type and metadata information.

## References

1. Amir, R. E. et al. Rett syndrome is caused by mutations in X-linked MECP2, encoding methyl-CpG-binding protein 2. Nat. Genet. 23, 185–188 (1999).

2. Zoghbi, H. Y. MeCP2 dysfunction in humans and mice. J. Child Neurol. 20, 736–740 (2005).

3. Sharifi, O. & Yasui, D. H. The Molecular Functions of MeCP2 in Rett Syndrome Pathology. Front. Genet. 12, (2021).

4. Yasui, D. H. et al. Mice with an isoform-ablating Mecp2exon 1 mutation recapitulate the neurologic deficits of Rett syndrome. Hum. Mol. Genet. 23, 2447–2458 (2014).

5. Guy, J., Hendrich, B., Holmes, M., Martin, J. E. & Bird, A. A mouse Mecp2-null mutation causes neurological symptoms that mimic rett syndrome. Nat. Genet. 27, 322–326 (2001).

6. Braunschweig, D., Simcox, T., Samaco, R. C. & LaSalle, J. M. X-chromosome inactivation ratios affect wild-type MeCP2 expression within mosaic Rett syndrome and Mecp2-/+ mouse brain. Hum. Mol. Genet. (2004). doi:10.1093/hmg/ddh142

7. Ballas, N., Lioy, D. T., Grunseich, C. & Mandel, G. Non-cell autonomous influence of MeCP2-deficient glia on neuronal dendritic morphology. Nat. Neurosci. (2009). doi:10.1038/nn.2275

8. Belichenko, P. V et al. Widespread Changes in Dendritic and Axonal Morphology in Mecp2-Mutant Mouse Models of Rett Syndrome: Evidence for Disruption of Neuronal Networks. J. Comp. Neurol. 514, 240–258 (2009).

9. Neier, K. et al. Sex disparate gut microbiome and metabolome perturbations precede disease progression in a mouse model of Rett syndrome. doi:10.1038/s42003-021-02915-3

10. Yasui, D. H. et al. Mice with an isoform-ablating Mecp2exon 1 mutation recapitulate the neurologic deficits of Rett syndrome. Hum. Mol. Genet. 23, 2447–2458 (2014).

11. Vogel Ciernia, A. et al. Early motor phenotype detection in a female mouse model of Rett syndrome is improved by cross-fostering. Hum. Mol. Genet. 26, 1839–1854 (2017).

12. tasic, B. et al. Shared and distinct transcriptomic cell types across neocortical areas. Nature (2018). doi:10.1038/s41586-018-0654-5

13. Mou, T., Deng, W., Gu, F., Pawitan, Y. & Nghia Vu, T. Reproducibility of Methods to Detect Differentially Expressed Genes from Single-Cell RNA Sequencing. doi:10.3389/fgene.2019.01331

14. Soneson, C. & Robinson, M. D. Bias, robustness and scalability in single-cell differential expression analysis. Nat. Methods 15, (2018).

15. Liu, Y. et al. iDESC: identifying differential expression in single-cell RNA sequencing data with multiple subjects. BMC Bioinformatics 24, (2023).

16. Renthal, W. et al. Characterization of human mosaic Rett syndrome brain tissue by single-nucleus RNA sequencing. Nat. Neurosci. (2018). doi:10.1038/s41593-018-0270-6

17. Morabito, S. et al. Single-nucleus chromatin accessibility and transcriptomic characterization of Alzheimer’s disease. Nat. Genet. doi:10.1038/s41588-021-00894-z

18. Morabito, S., Reese, F., Rahimzadeh, N., Miyoshi, E. & Swarup, V. hdWGCNA identifies co-expression networks in high-dimensional transcriptomics data. doi:10.1016/j.crmeth.2023.100498

19. Langfelder, P. & Horvath, S. WGCNA: an R package for weighted correlation network analysis. (2008). doi:10.1186/1471-2105-9-559

20. Bakken, T. E. et al. Comparative cellular analysis of motor cortex in human, marmoset and mouse. doi:10.1038/s41586-021-03465-8

21. Tudor, M., Akbarian, S., Chen, R. Z. & Jaenisch, R. Transcriptional profiling of a mouse model for Rett syndrome reveals subtle transcriptional changes in the brain. Proc. Natl. Acad. Sci. U. S. A. 99, (2002).

22. Nuber, U. A. et al. Up-regulation of glucocorticoid-regulated genes in a mouse model of Rett syndrome. Hum. Mol. Genet. 14, (2005).

23. Kriaucionis, S. et al. Gene Expression Analysis Exposes Mitochondrial Abnormalities in a Mouse Model of Rett Syndrome. Mol. Cell. Biol. 26, (2006).

24. Urdinguio, R. G. et al. Mecp2-null mice provide new neuronal targets for rett syndrome. PLoS One 3, (2008).

25. Chahrour, M. et al. MeCP2, a key contributor to neurological disease, activates and represses transcription. Science (80-.). (2008). doi:10.1126/science.1153252

26. Ben-Shachar, S., Chahrour, M., Thaller, C., Shaw, C. A. & Zoghbi, H. Y. Mouse models of MeCP2 disorders share gene expression changes in the cerebellum and hypothalamus. Hum. Mol. Genet. 18, (2009).

27. Samaco, R. C. et al. Crh and Oprm1 mediate anxiety-related behavior and social approach in a mouse model of MECP2 duplication syndrome. Nat. Genet. 44, (2012).

28. Sugino, K. et al. Cell-type-specific repression by methyl-CpG-binding protein 2 is biased toward long genes. J. Neurosci. 34, (2014).

29. Osenberg, S. et al. Activity-dependent aberrations in gene expression and alternative splicing in a mouse model of Rett syndrome. Proc. Natl. Acad. Sci. U. S. A. 115, (2018).

30. Zilionis, R. et al. Single-cell barcoding and sequencing using droplet microfluidics. Nat. Publ. Gr. (2016). doi:10.1038/nprot.2016.154

31. Zhang, X. et al. Comparative Analysis of Droplet-Based Ultra-High-Throughput Single-Cell RNA-Seq Systems Molecular Cell Comparative Analysis of Droplet-Based Ultra-High-Throughput Single-Cell RNA-Seq Systems. Mol. Cell 73, 130–142 (2019).

32. Deng, J. V. et al. MeCP2 in the nucleus accumbens contributes to neural and behavioral responses to psychostimulants. Nat. Neurosci. 13, (2010).

33. Su, D., Cha, Y. M. & West, A. E. Mutation of Mecp2 alters transcriptional regulation of select immediate-early genes. Epigenetics 7, (2012).

34. Zhang, W., Peterson, M., Beyer, B., Frankel, W. N. & Zhang, Z. W. Loss of MeCP2 from forebrain excitatory neurons leads to cortical hyperexcitation and seizures. J. Neurosci. 34, 2754–2763 (2014).

35. Sun, J. et al. Mutations in the transcriptional regulator MeCP2 severely impact key cellular and molecular signatures of human astrocytes during maturation. Cell Rep. 42, 111942 (2023).

36. Neul, J. L. et al. nature medicine Trofinetide for the treatment of Rett syndrome: a randomized phase 3 study. Nat. Med. | 29, 1468–1475 (2023).

37. Puche, J. E. & Castilla-Cortázar, I. Human conditions of insulin-like growth factor-I (IGF-I) deficiency. Journal of Translational Medicine 10, (2012).

38. Yao, Z. et al. A taxonomy of transcriptomic cell types across the isocortex and hippocampal formation. (2021). doi:10.1016/j.cell.2021.04.021

39. tasic, B. et al. Shared and distinct transcriptomic cell types across neocortical areas. Nature (2018). doi:10.1038/s41586-018-0654-5

