## Supplementary Figure 1-12 for "Sex-specific single cell-level transcriptomic signatures of Rett syndrome disease progression"

**a**

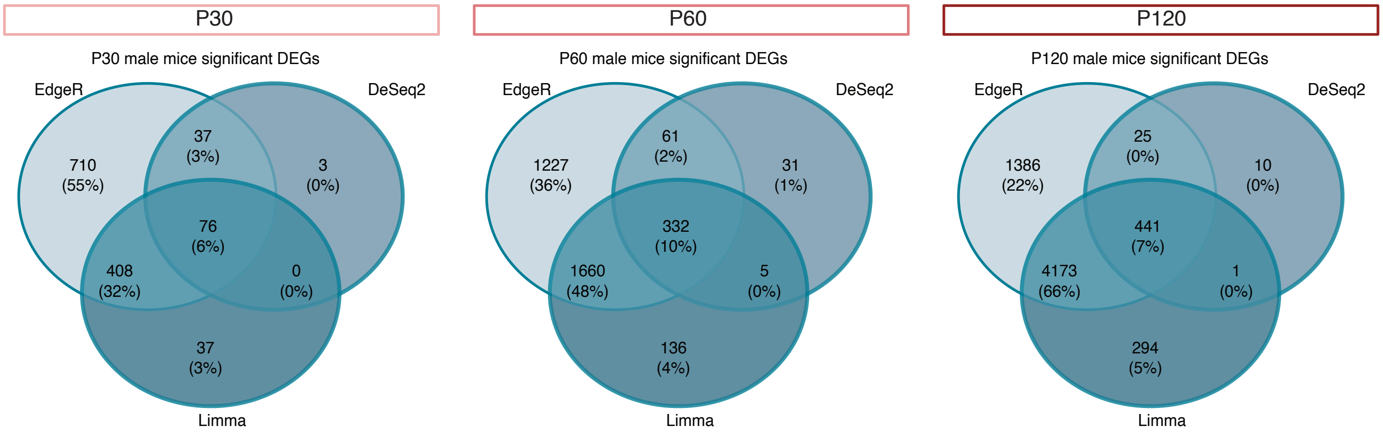


**Supplemental Figure 1. Identification of significant DEGs using Limma, EdgeR and DESeq2 produced few common genes from male sn-RNA seq data. a** Venn diagrams of the three DEG methods at P30, P60 and P120

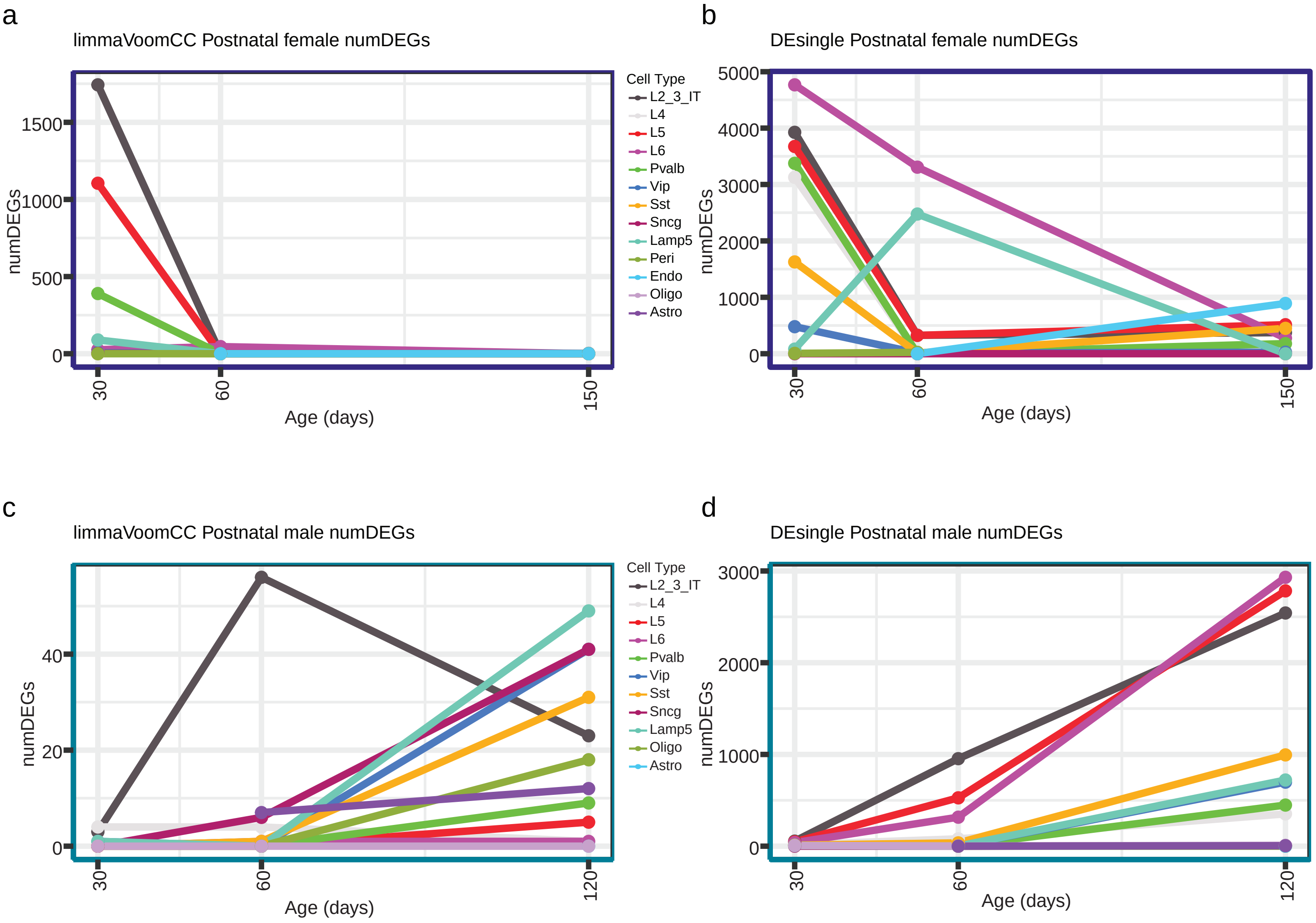


Supplemental Figure 2. Line graph of the number of post-natal DEGs identified using DESingle from *0HFS H* female and *0HFS H* cortices. a,b VKRZ IHPDOH '(*V DGMXVWHG S WLPH XVLQJ OLPPD9RRP&& DQG '(VLQJOH UHVSHFWLYHO\ c,d VKR RYHU WLPH XVLQJ OLPPD9RRP&& \DQG '(VLQJOH UHVSHFWLYHO

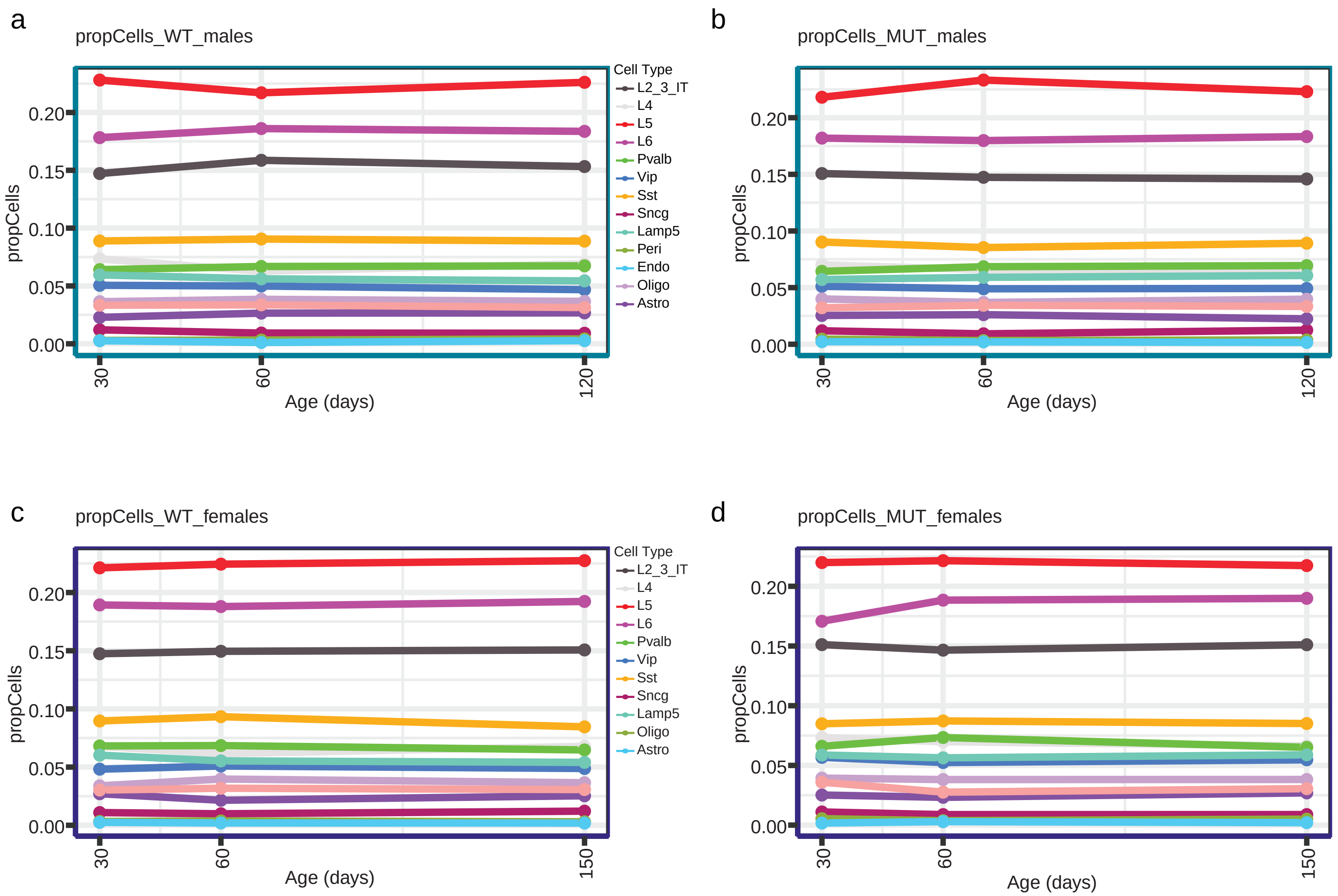


Supplemental Figure 3. 0HFS H mutant cortices have normal proportions of cortical cell types. a,b show line graphs of cell proportions in WT males and MUT males respectively over time. c,d show line graphs of cell proportions in WT females and MUT females respectively over time.

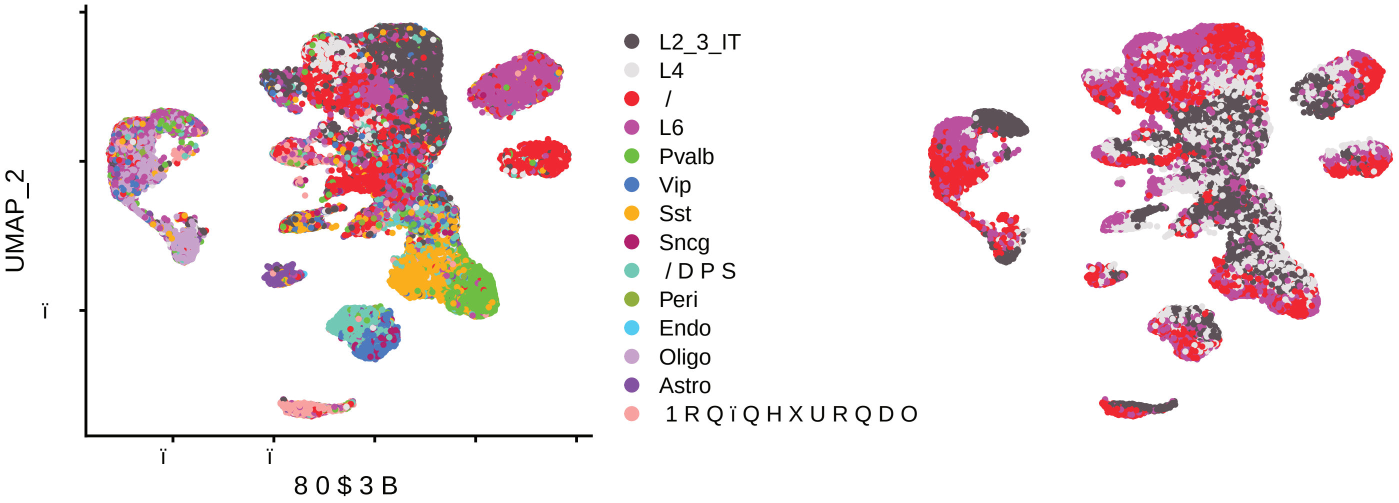

UMAP_2
ï
ï
ï
8 0 $ 3 B
L2_3_IT
L4
/
L6
Pvalb
Vip
Sst
Sncg
/ D P S
Peri
Endo
Oligo
Astro
1 R Q ï Q H X U R Q D O
